# Circulating tumor extracellular vesicles to monitor metastatic prostate cancer genomics and transcriptomic evolution

**DOI:** 10.1101/2023.04.14.536404

**Authors:** Irene Casanova-Salas, Sarai Cordoba-Terreros, Daniel Aguilar, Laura Agundez, Julian Brandariz, Nicolas Herranz, Macarena Gonzalez, Rafael Morales-Barrera, Alexandre Sierra, Mario Soriano-Navarro, Pablo Cresta, Sara Simonetti, Gonçalo Rodrigues, Sara Arce-Gallego, Luisa Delgado-Soriano, Irene Agustí, Elena Castellano-Sanz, Richard Mast, Matias de Albert, Ana Celma, Anna Santamaria, Lucila Gonzalez, Natalia Castro, Maria del Mar Suanes, Javier Hernández-Losa, Lara Nonell, Hector Peinado, Joan Carles, Joaquin Mateo

**Affiliations:** Vall d’Hebron Institute of Oncology (VHIO), Barcelona, Spain; Vall d’Hebron Institute of Research (VHIR), Barcelona, Spain; Vall d’Hebron University Hospital, Barcelona, Spain; Principe Felipe Research Institute (CIPF), Valencia, Spain; Microenvironment and Metastasis Laboratory, Molecular Oncology Program, Spanish National Cancer Research Center (CNIO), Madrid, Spain

**Keywords:** prostate cancer, biomarkers, liquid biopsy, extracellular vesicles, transcriptomics.

## Abstract

Extracellular vesicles (EVs) secreted by tumors are abundant in plasma, but their potential for interrogating the molecular features of tumors through multi-omic profiling remains widely unexplored. Genomic and transcriptomic profiling of circulating EV-DNA and EV-RNA isolated from a range of *in-vitro* and *in-vivo* models of metastatic prostate cancer (mPC) revealed a high contribution of tumor material to EV-loaded DNA/RNA. Findings were validated in a cohort of longitudinal plasma samples collected from mPC patients during androgen receptor signaling inhibitor (ARSI) therapy. EV-DNA genomic features recapitulated matched-patient biopsies and associated with clinical progression. We developed a novel approach to enable the transcriptomic profiling of EV-RNA (RExCuE). We report how the transcriptomic profile in mPC EV-RNA is enriched for tumor-associated transcripts when compared to same patient blood RNA and healthy individuals EV-RNA, and reflect early on-therapy tumor adaptation changes. Altogether, we show that EV profiling enables longitudinal transcriptomic and genomic profiling of mPC in liquid biopsy.

## Introduction

Metastatic prostate cancer (mPC) is a lethal disease; while second-generation AR signaling inhibitors (ARSI) have improved the survival of patients with mPC (1) responses are largely heterogeneous, and tumors eventually develop resistance to therapy (2,3). Developing biomarkers that can guide clinical decisions in mPC is key for a more precise patient care.

Several studies have identified transcriptional plasticity as a key mechanism underlying ARSI resistance (3–6). Longitudinal transcriptomic profiling of tumor samples throughout treatment could inform, or potentially anticipate, drug resistance mechanisms and guide subsequent therapies. However, obtaining repeatedly suitable tumor material for molecular profiling in clinical practice remains challenging.

Liquid biopsy represents an emerging tool for molecular characterization and longitudinal disease monitoring. Next-generation sequencing (NGS) of circulating tumor DNA (ctDNA) has proven valuable for clinical management and therapeutic selection in mPC (7,8). In contrast, direct interrogation of tumor transcriptomic features from plasma cell-free RNA (cfRNA) has been unsuccessful because of the rapid degradation of RNA when it is not protected by the cell membrane; while RNA analysis of circulating tumor cells (CTCs) (9) has not been translated into routine clinical practice because of the technical challenges associated with CTC isolation. Tumors also secrete extracellular vesicles (EVs) into their bloodstream. EVs are characterized by a lipid bilayer and a typical size between 50 nm and 1 µm; they contain proteins, lipids, metabolites, RNA (mRNA, lncRNA, miRNAs, etc.), and DNA as cargo. EVs produced by tumor cells are key mediators of cancer cell communication, and the protein cargo composition of EVs plays a role in tumor progression, immune regulation, and metastasis (10,11). However, the potential of tumor EVs as a source of clinically relevant DNA and RNA biomarkers remains largely unexplored. We hypothesized that EVs shed by prostate cancer (PC) cells would enable tumor genomic and transcriptomic characterization, opening avenues for biomarkers that can study tumor adaptation at the time of therapy response and resistance.

In this study, we combined DNA and RNA NGS to analyze the DNA and protein-coding RNA content of circulating EVs (EV-DNA and EV-RNA) in mPC, including clinical samples from patients undergoing treatment with ARSI (abiraterone acetate or enzalutamide). We propose that our approach for EV-RNA sequencing offers a tool to identify the mechanisms of resistance and monitor tumor-associated changes upon therapy intervention.

## Results

### Characterization of circulating EV-DNA in prostate cancer patients

EVs were first isolated using ultracentrifugation from the conditioned media of PC cell line models (LNCaP, C4-2, 22Rv1, and PC3) and from plasma from 22Rv1-derived tumor-bearing mouse xenografts. The presence of tetraspanins (CD9 and CD81) and TSG101 suggested an enrichment of small EV (sEV) (**Supplementary Figure 1A**). Transmission electron microscopy (TEM) analysis confirmed the presence of cup-shaped structures with a lipid bilayer and diameter of 50-150 nm, corresponding to the morphology and size characteristics of sEVs (**Supplementary Figure 1B**). This was confirmed by Nanosight analysis (NTA), which revealed small vesicles with a median diameter of 124–141 nm (**Supplementary Figure 1C**). Purity of the isolated sEV was assessed by further separation of the sEVs through iodixanol density gradient (**Supplementary Figure 1D**). Small EV markers CD81 and TSG101 were present in fractions F7-F10. We also confirmed that DNA concentration was higher in sEVs than in larger vesicles (12K) and found that, particularly, F8 showed the highest DNA concentration. Low-pass whole genome sequencing (lpWGS) of EV-DNA revealed representation of the entire tumor genome in circulating EVs: copy number alteration (CNA) profiles for EV-DNA were highly concordant (r 0.7-0.99 across comparisons; all P<0.0001) with those of their matching tumor cells *in vitro* and in plasma from tumor xenografts (**Supplementary Figure 1E**).

Together, these data prompted us to evaluate EV-DNA in mPC patient samples. We first characterized EVs isolated from plasma samples of a retrospective cohort of 22 patients with mPC. We confirmed the expression of sEV protein markers (TSG101, CD9, and CD63) (**Supplementary Figure 2A)**, TEM analysis verified the presence of sEVs (**Supplementary Figure 2B**), and NTA showed vesicles with a median diameter of 102-133 nm (**Supplementary Figure 2C**).

Next, we isolated both EV-DNA and cfDNA from plasma samples in a further cohort of 35 castration-resistant mPC patients with plasma collected at different time points during ARSI treatment (baseline, BL; on treatment, On-ttx; and at progression, PD) (**Figure 1A** and **Table 1**). Over 80% of EV-DNA in plasma was found to be DNAse-protected inside these vesicles, whereas cfDNA was completely degraded after DNAse digestion, confirming these are two different biological entities (**Figure 1B**). For all subsequent experiments, we performed DNAse digestion of all EV preparations prior to DNA isolation to ensure that EV-DNA did not contain any cfDNA. EV-DNA had a different dsDNA size distribution than cfDNA in mPC patient samples: cfDNA was enriched for smaller DNA fragments (100-200bp) whereas EV-DNA was enriched for larger fragments (>1000bp) (P<0.001) (**Figure 1C**).

**Figure 1—.**
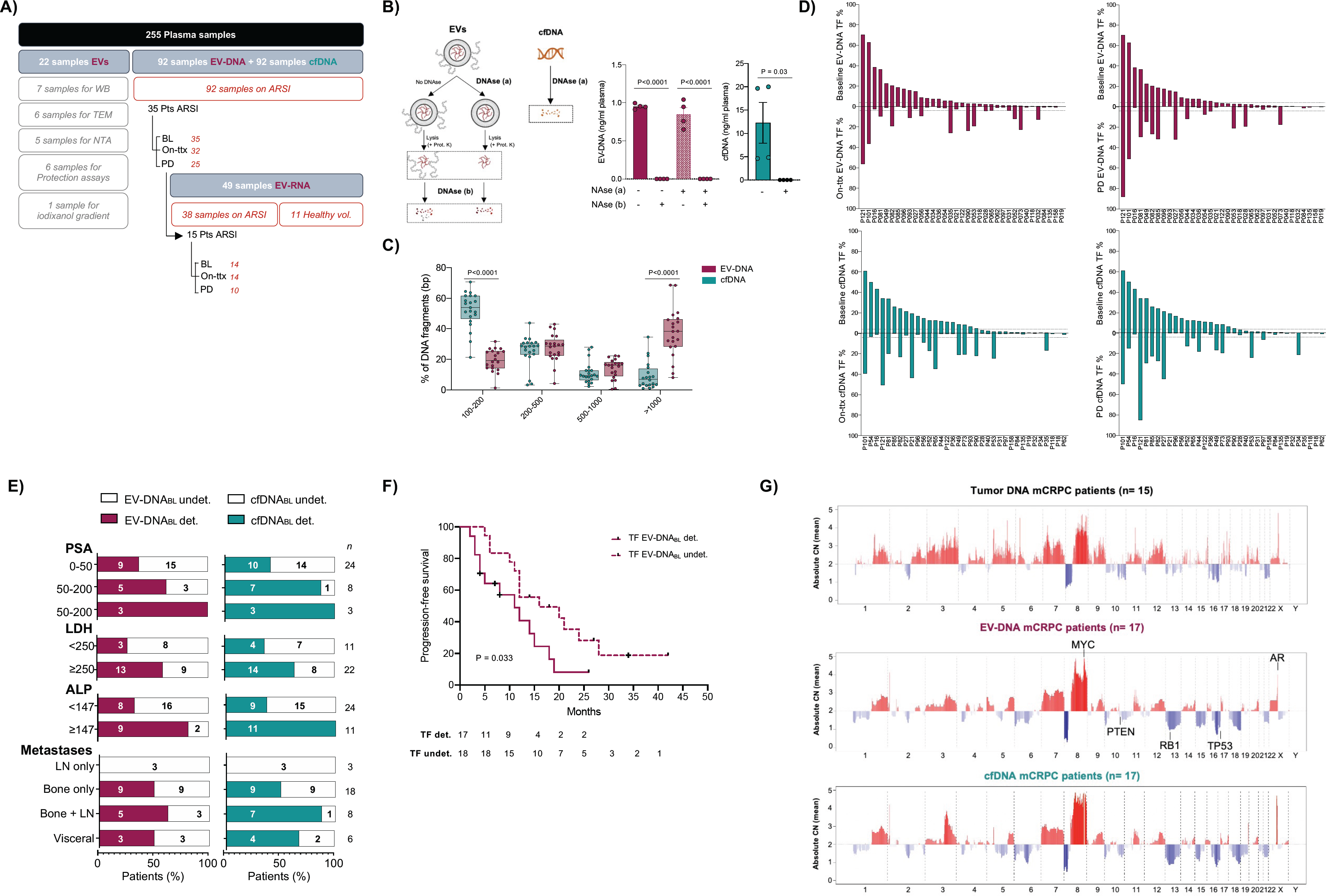
Copy number alteration (CNA) profiling of circulating EV-DNA in mPC confirms tumor origin and identifies patients at higher risk of progression. **a)** Representative scheme of the study design and sample disposition b) Enzymatic protection assay experiment and graph bar showing tapestation quantification of dsDNA concentration (ng/mL plasma) from untreated and DNAse I-treated samples from plasma-derived EV-DNA and cfDNA. **c)** Size-distribution of dsDNA fragments in plasma-derived EV-DNA and cfDNA from mPC patients. Boxplot showing the percentage of fragments of different sizes found in each patient’s EV-DNA and cfDNA collected (n=35 patients). **d)** Mirrored bar plot showing same-patient tumor fraction (TF) in plasma-derived EV-DNA and cfDNA at baseline vs 4-weeks after treatment (On-ttx) or at baseline vs progression (PD). **e)** Fraction of patients with detectable (det.) or undetectable (undet.) EV-DNA and cfDNA tumor fraction at baseline (BL) according to different clinical variables. **f)** Kaplan-Meier curves for progression-free survival on ARSI therapy according to tumor fraction in EV-DNA at baseline (TF EV-DNA_BL_ detectable vs undetectable, using a 3% TF threshold based on ichorCNA). P-values shown were determined using the log-rank test. **g)** Representation of the average whole genome CNA profile calculated for matching mCRPC tumor biopsies (n=15) and plasma-derived EV-DNA and cfDNA at baseline with detectable TF (n=17). Amplifications are depicted in red and deletions in blue. Key driver genes for mPC are highlighted in EV-DNA CNA profile.

**Table 1.** Patient characteristics at baseline.

|  | Prospective cohort (n = 35) |
| --- | --- |
| **Age** | 74,3 (50-91) |
| **PSA (ng/ml)** | 82.2 (2.3-1066) |
| **LDH (UI/L)** | 333.9 (109-770) |
| **ALP (UI/L)** | 193.1 (50-1295) |
| **Sites of metastasis** |  |
| Bone only | 18 (51,4%) |
| Lymph node and bone | 8 (22,8%) |
| Lymph node only | 3 (8,6%) |
| Visceral | 6 (17,2%) |
| **Current treatment** |  |
| Abiraterone | 15 (42.8%) |
| Enzalutamide | 20 (57.2%) |
| **Prior treatment** |  |
| ADT | 35 (100%) |
| Enzalutamide | 1 (2.8%) |
| Docetaxel | 3 (8.6%) |
| Cabazitaxel | 1 (2.8%) |

Concentration of DNA loaded in EVs (mean:0.61 ng/ml plasma; 95%CI 0.46-0.76) was almost 50-fold lower than cfDNA (mean:23.7 ng/ml plasma; 95%CI 14-33.4) (**Supplementary Figure 3A)**. Tumor fraction for each analyte, calculated from lpWGS, was similar between EV-DNA and cfDNA, with a TF range of 0.3-88.3% for EV-DNA and 0.2-85% for cfDNA (**Figure 1D** and **Supplementary Figure 3B**). A detectable tumor fraction (defined as TF>3% based on ichorCNA) was found in 17/35 EV-DNA and 20/35 cfDNA treatment-baseline samples. Among patients with detectable EV-DNA TF at baseline, in 7/17 EV-DNA TF remained detectable on treatment, and 10/17 at progression. Whereas for patients with cfDNA TF detected at baseline, 11/20 had detectable TF on treatment and 12/20 at progression (**Figure 1D**). Interestingly, there were some patients and time points where TF was higher in EV-DNA vs. cfDNA or higher in cfDNA vs. EV-DNA (**Figure 1D** and **Supplementary Figure 3B-C, Supplementary Table S1**), highlighting the potential complementarity of the dual cfDNA/EV-DNA analysis.

A detectable TF in EV-DNA was associated with clinical markers of tumor burden, including prostate-specific antigen (PSA), lactate dehydrogenase (LDH), alkaline phosphatase (ALP) and presence of bone metastases (**Figure 1E**). EV-DNA tumor fraction was correlated with patient outcome: an EV-DNA TF>3% at baseline associated with significantly shorter time to progression (HR = 2.29; 95% CI, 1.05-5.12; log-rank test P = 0.033; **Figure 1F, Supplementary Figure 3D**). We studied the CNA profiles by lpWGS in matching mPC tumor biopsies, plasma EV-DNA and cfDNA at baseline. We confirmed the representation of the entire genome and the high correlation between plasma-EV-DNA (r = 0.73, P<0.0001), and cfDNA (r = 0.75, P<0.0001) with patient-matched tumor tissue biopsies (TBx DNA) (**Figure 1G**). These findings confirm the presence of tumor DNA cargo in circulating EVs and supports the use of sEVs obtained from liquid biopsies to interrogate the genomic features of tumors.

### Development of a pipeline for EV-mRNA analysis

To investigate the transcriptomic information encapsulated in EVs, we characterized EV-RNA in plasma from mPC patients in our cohort that had a detectable EV-DNA TF, aiming to prioritize those cases likely to have more tumor material in circulation. EVs isolated for RNA analysis (column-based isolation method, see Methods) had a median size of 180-200 nm (**Supplementary Figure 4A**). These vesicles expressed traditional sEV protein markers and exhibited a characteristic cup-shaped morphology (**Supplementary Figure 4B**). Purity of sEVs isolated for EV-RNA studies was confirmed by further separation of sEVs through iodixanol density gradient (**Supplementary Figure 4C**). We found that sEV markers CD9 and TSG101 were present in fraction F8, and fraction F7 and F8 had the highest RNA concentration.

EV-RNA molecules isolated from mPC patient plasma samples showed significant representation of RNA molecules with a size between 50-200nt (P<0.0001) (**Figure 2A**). We treated plasma EVs with RNAse pre-EV lysis, confirming that most RNA isolated using our methodology was RNAse-protected inside these vesicles (**Supplementary Figure 4D**).

**Figure 2—.**
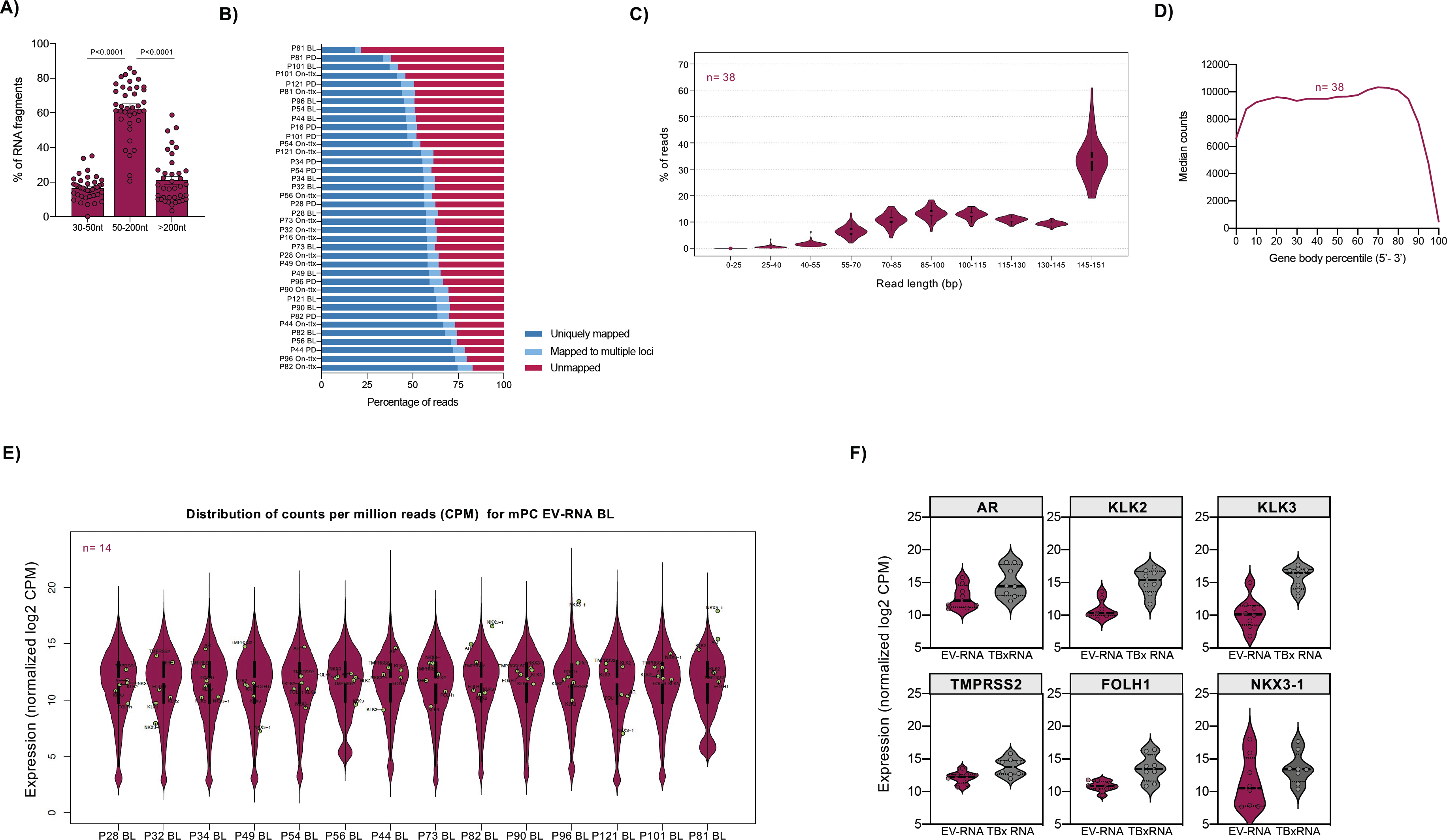
RExCuE development for mRNA analysis from circulating EVs. **a)** Size-distribution of RNA fragments in circulating EV-RNA collected from mPC patient plasma (n=38). Scatter plot showing the percentage of fragments of different sizes found in EV-RNA. P-value<0.0001. **b)** Graph bar showing the summary mapping statistics for longitudinally collected plasma EV-RNA samples (n=38) following RExCuE pipeline. **c)** Length distribution (mean ± SEM) of uniquely mapped reads from longitudinal plasma EV-RNA (n=38). **d)** Gene body coverage on average for all genes expressed in plasma EV-RNA at baseline. X-axis represents the positions within the gene (on average), while y-axis represents the average read count for each position. **e)** Violin plots showing normalized gene expression distribution (mean ± SEM) for mPC plasma EV-RNA samples at baseline. Key PC genes are highlighted in green within the average distribution of counts. **f)** Violin plots showing normalized gene expression distribution (mean ± SEM) for a selected group of prostate-specific genes in baseline EV-RNA and matching patients tumor biopsy RNA (TBx RNA)

We developed a methodology to specifically capture protein-coding RNA from these small RNA molecules, and to prepare libraries for assessing m**R**NA **Ex**pression in **C**irculating **E**Vs (RExCuE libraries). Despite the size distribution of our starting RNA material, accurate alignment (average 60.7% mapped reads) was obtained for almost all plasma EV-RNA samples using RExCuE (**Figure 2B**). We sequenced an average of 50 million reads and obtained an average of 29 million uniquely mapped reads for further downstream analysis. We obtained a homogenous read length distribution, with the majority of uniquely mapped reads with a size between 145-151 bp (**Figure 2C**). We further confirmed that these uniquely mapped reads were able to uniformly cover the whole transcript length of all detected genes (**Figure 2D**).

RExCuE allowed us to generate expression data for over 30,000 genes in plasma EV-RNA with a gene expression distribution of 0.9-24 counts per million reads (CPM) (**Figure 2E**). Over 70% mapped transcripts corresponded to protein-coding genes, with smaller representation of lncRNA (20%) and miRNAs (5%) (**Supplementary Figure 4E**).

Next, we confirmed the presence of PC-associated transcripts: AR, KLK2, KLK3, TMPRSS2, FOLH1 and NKX3-1 in all plasma EV-RNA samples, as well as in matching tumor biopsies RNA (TBx RNA) (**Figure 2E-F**).

In summary, RExCuE delivers plasma EV-RNAseq data of good quality that provides transcriptomic information in circulation of relevant genes in mPC.

### Analysis of EV-RNA in metastatic prostate cancer patients compared to plasma from healthy volunteers and patient-matched PBMCs

We compared the transcriptome of plasma EV-RNA from mPC patients (n=14 pre-treatment samples) and age-matched healthy volunteers (HV, n=7). The resulting principal component analysis (PCA) showed that samples positioned differently according to their gene expression depending on their patient vs HV origin (**Figure 3A)**. Differential expression (DE) analysis identified >200 genes significantly up or down-regulated in patients’ samples (**Figure 3B)**. Interestingly, genes highly up-regulated in EV-RNA from mPC patients included several genes associated with keratins (KRTAP5-2, KRTAP17-1, KRTAP4-5, KRTAP5-5) and other tumor-related processes. GO analysis revealed that those genes up-regulated in mPC EV-RNA were enriched for biological processes such as tissue development, epithelium development and differentiation and keratinization (**Supplementary Figure 5A**).

**Figure 3—.**
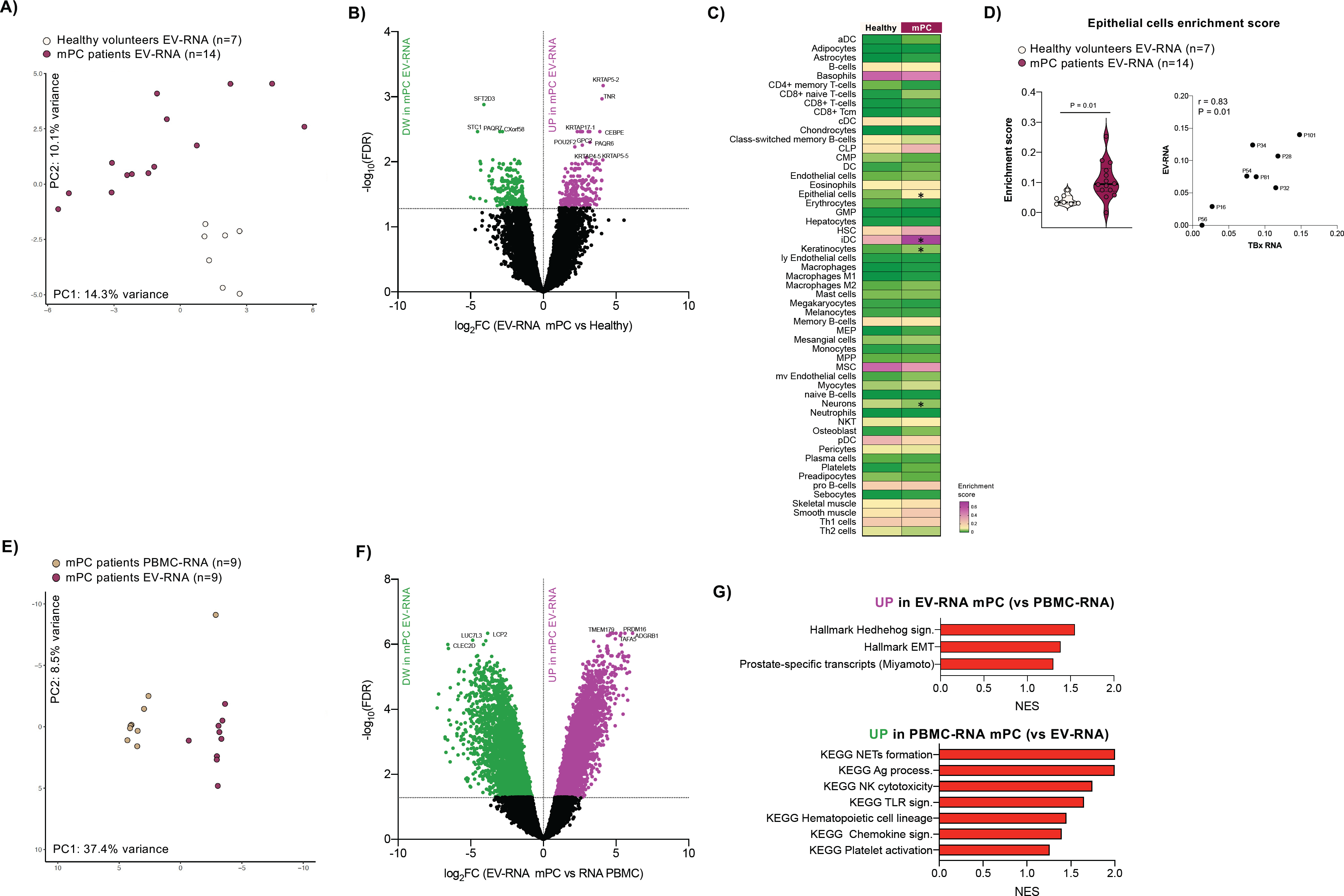
EV-RNA gene expression profiling identifies tumor-associated transcripts in mPC EV-RNA. **a)** Principal component analysis (PCA) of EV-RNA transcriptomes from healthy volunteers (white, n=7) and mPC patients at baseline (magenta, n=14). **b)** Volcano plot showing DEG in EV-RNA from mPC patients at baseline vs. healthy volunteers’ EV-RNA. Statistically significant (FDR<0.05) up-and down-regulated genes are depicted in purple and green respectively. Top up and down regulated genes in mPC EV-RNA samples are highlighted. **c)** Cell type enrichment analysis in EV-RNAseq data from healthy volunteers (n=7) and mPC patients’ plasma (n=14). Average enrichment scores for healthy and mPC EV-RNA are shown in the heatmap. Asterisk shows statistically different cell types in mPC vs. healthy EV-RNA (P<0.05). **d)** Left, violin plots showing distribution of individual enrichment scores for the epithelial subtype in healthy volunteers and mPC patients’ EV-RNA. Right, similarity in the calculated epithelial enrichment score between EV-RNA and matching tumor biopsy RNA (TBx RNA) is given by Pearson correlation score (r) and P-value. **e)** Principal component analysis (PCA) of transcriptomes from mPC patients’ EV-RNA (magenta, n=9) and matching PBMCs (brown, n=9). **f)** Volcano plot showing DEG in paired EV-RNA vs. PBMCs’ RNA from mPC patients. Statistically significant (FDR<0.05) up-and down-regulated genes are depicted in purple and green respectively. Top up and down regulated genes in mPC EV-RNA samples are highlighted. **g)** Gene-set enrichment analysis (GSEA) performed on genes up-regulated in EV-RNA vs. PBMCs’ RNA (above) or up-regulated in PBMCs’ RNA vs. EV-RNA. FDR<0.25 was used for significance. Normalized enrichment score (NES).

Cell deconvolution pipelines, using xCell (12), were applied to our EV-RNAseq data from healthy volunteers and mPC patients to further understand the contribution of different cell types to the transcriptomic signal detected in circulation (**Figure 3C**). As expected, average scores of cell types in plasma EV-RNA from both healthy volunteers and mPC patients showed enrichment of blood-related cell types such as hematopoietic stem cells, lymphocytes, and myeloid cells. However, EV-RNA from mPC patients showed a significantly higher enrichment for cell types commonly enriched in carcinomas (12) such as epithelial cells (P=0.01) and keratinocytes (P=0.05), further supporting the prostate cancer origin of this material (**Supplementary Figure 5B-C**). This enrichment for epithelial cells observed in mPC EV-RNA was further confirmed in matching tumor biopsy RNA (TBx RNA) (r=0.83, P=0.01) (**Figure 3D**).

Different blood cells shed EVs into circulation, contributing significantly to the overall cargo of the EV pool in circulation (13,14). Therefore, as additional control, we pursued RNAseq from same-patient PBMCs, isolated from the same blood sample used for EV-RNA isolation. PCA results clearly separated samples according to origin of the sample (EV vs PBMC) (**Figure 3E**). DE analysis identified over 10000 genes significantly differentially expressed between EV-RNA and PBMC-RNA (**Figure 3F**), with an enrichment for PC associated transcripts (15), as well as tumor-associated pathways (such as hedhegog signaling and epithelial to mesenchymal transition) in EV-RNA compared to same patient’s PBMCs. While we found enrichment of pathways related with blood lineage and immune system in PBMC-RNA when compared to plasma EV-RNA (**Figure 3G**).

Overall, these results indicate that the transcriptome profile found in plasma EV-RNA from mPC patients is distinct to plasma EV-RNA from healthy volunteers and PBMC-RNA, and is enriched for RNA of tumoral origin.

### Transcriptomic signatures in EV-RNA upon AR inhibition

To investigate the potential of EV-RNA as response and resistance biomarker, we studied changes in circulating EV-RNA upon treatment with ARSI enzalutamide in PC models, both *in vitro* (LNCaP and C4-2 cells were treated with vehicle or enzalutamide for 24h) and *in vivo* (LNCaP and C4-2 xenografts were randomized to either vehicle or enzalutamide treatment, **Figure 4A-B**). As expected, LNCaP xenograft tumors were highly sensitive to enzalutamide, while C4-2 castrated mice showed a milder reduction in tumor growth.

**Figure 4—.**
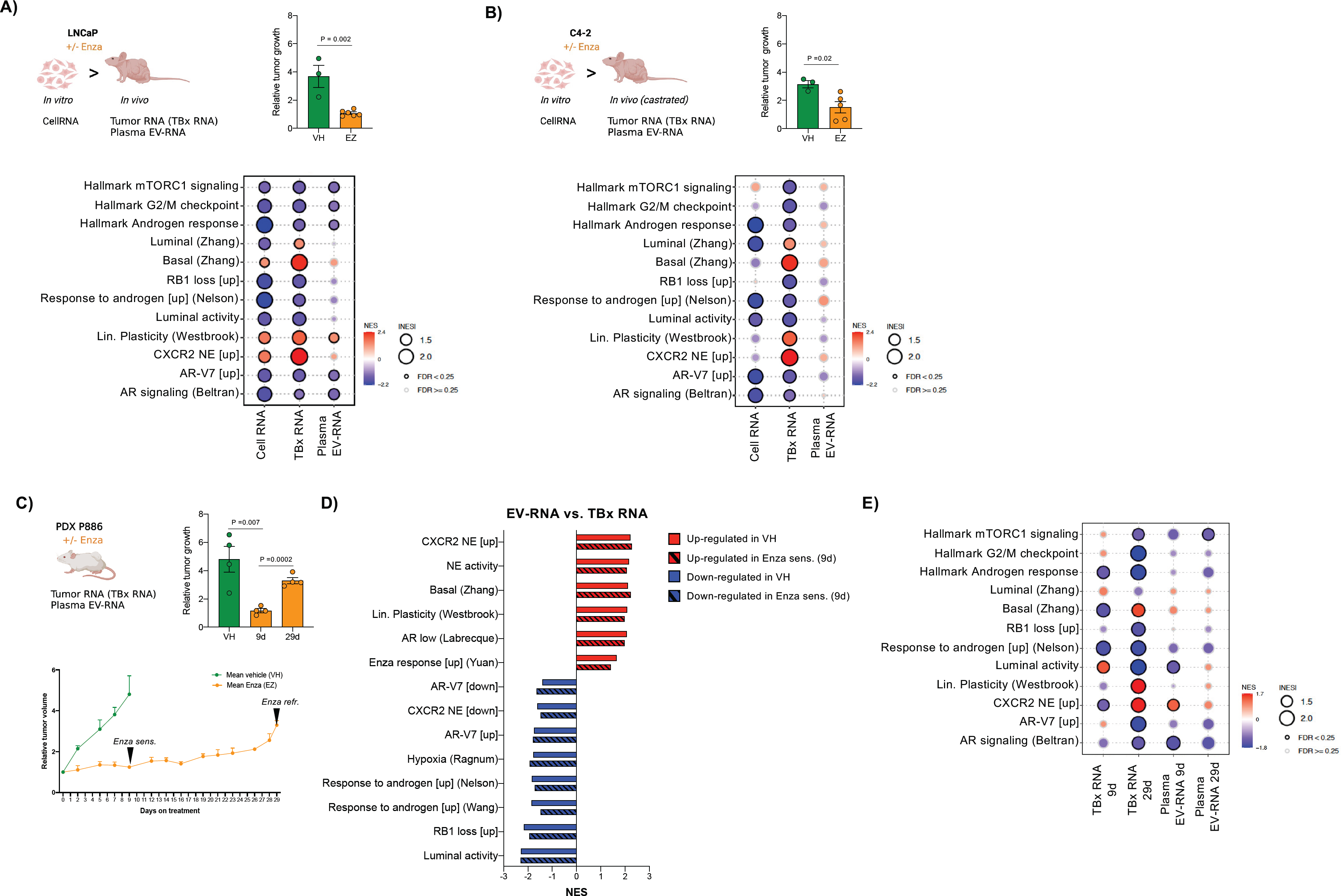
EV-RNA gene expression profiling identifies transcriptomic signatures of response and resistance to enzalutamide. **a)** Left, scheme of the samples used for LNCaP *in vitro* and *in vivo* analysis. Right, representation of the average relative tumor growth in LNCaP xenografts treated with enzalutamide (orange; n=6) or vehicle (green; n=3). Bubble plot shows up and down-regulated gene signatures within MSigDB hallmark and literature curated gene sets for enzalutamide versus vehicle in LNCaP cells *in vitro* (CellRNA), *in vivo* (TBx RNA), and in xenografts’ plasma EV-RNA (Plasma EV-RNA). **b)** Left, scheme of the samples used for C4-2 *in vitro* and *in vivo* analysis. Right, representation of the average relative tumor growth in C4-2 castrated xenografts treated with enzalutamide (orange; n=5) or vehicle (green; n=3). Bubble plot shows up and down-regulated gene signatures within MSigDB hallmark and literature curated gene sets for enzalutamide versus vehicle in C4-2 cells *in vitro* (CellRNA), *in vivo* (TBx RNA), and in xenografts’ plasma EV-RNA (Plasma EV-RNA). **c)** Scheme of the samples used for PDX P886 analysis., Graph bar represents the average relative tumor growth in PDX treated with enzalutamide for 9 or 29 days (orange; n= 4) or vehicle (green; n=2). Below, relative tumor growth curves show enzalutamide sensitive (Enza sens; 9 days) and enzalutamide refractory periods. PDX P886 (Enza refr; 9 days). **d)** Bubble plot showing the top most significant gene set enrichment scores in EV-RNA versus Tumor RNA from vehicle and enzalutamide treated (9 days) PDXs. **e)** Bubble plot shows up and down-regulated gene signatures within MSigDB hallmark and literature curated gene sets for tumors (TBx RNA) and plasma-derived EV-RNA (Plasma EV-RNA) after 9 and 29 days of treatment with enzalutamide. FDR < 0.25 was used for significance in all gene-set enrichment analysis (GSEA). Normalized enrichment score (NES).

We performed RNAseq using our RExCuE approach on RNA from *in vitro* tumor cells (CellRNA), *in-vivo* tumor samples RNA (TBx RNA) and EV-RNA isolated from LNCaP and C4-2 xenografts plasma. Gene set enrichment analysis (GSEA) of LnCaP EV-RNA showed downregulation of metabolism-related genes (Hallmark mTORC1 signaling) and cell proliferation signatures (Hallmark G2/M checkpoint), a decrease in AR signaling related signatures and an increase in several signatures related to basal-like and neuroendocrine differentiation were observed upon enzalutamide therapy, parallel to the observation in both in *in-vitro* and *in-vivo* LnCaP tumor samples (**Figure 4A**). We also observed a decrease, albeit milder, in the expression of several AR-related signatures in C4-2 EV-RNA upon treatment with enzalutamide, as observed in C4-2 CellRNA and TBx RNA. Furthermore, basal-like and neuroendocrine signatures showed increased expression both in C4-2 TBxRNA and plasma EV-RNA (**Figure 4B**).

Next, we analyzed plasma EV-RNA from patient-derived xenograft (PDX) P886, a PDX model that presents a short-lasting response to enzalutamide. PDX-P886 was derived in our laboratory from a prostate biopsy of a patient with metastatic *de novo* PC presenting with bone metastasis, Gleason score 5+4 adenocarcinoma, and PSA level at diagnosis of 46 ng/ml. The patient experienced a short-lasting response to androgen deprivation therapy (ADT), with progression to castration-resistance after 6 months of ADT. PDX P886 were randomized to vehicle or enzalutamide: similar to the clinical observation, PDX P886 initially responded to enzalutamide (Enza sensitive period), and rapidly relapsed after a month (Enza refractory period) (**Figure 4C**). Intratumor heterogeneity for AR expression was noted at baseline, with an enrichment for AR-negative cells upon enzalutamide progression **(Supplementary Figure 6A)**. We collected tumors (TBx RNA) and plasma (EV-RNA) at day 9 (Enza sensitive) and 29 (Enza refractory) after enzalutamide/vehicle treatment. We confirmed the presence of tumor-derived EVs by analyzing the CNA profile of plasma EV-DNA which resemble the CNA profile from PDX P886 tumor (TBx) (r=0.92, P<0.001) (**Supplementary Figure 6B**). EV-RNA isolated from PDX P886 plasma was enriched for 50-200nt RNA fragments, similar to our observation in patient samples **(Supplementary Figure 6C)**.

FGSEA of EV-RNA compared to matching tumor RNA, both in vehicle and enzalutamide-treated mice, confirmed an enrichment for basal-like signatures and a relative lower expression of AR-associated signatures in EV-RNA, suggesting different disease subclones may be differentially represented in circulating material **(Figure 4D)**. EV-RNA after 9 or 29 days of treatment showed significant down-regulation of AR signatures, as observed in the PDX tumor samples. Interestingly, plasma EV-RNA at day 9 post-enzalutamide already showed an enrichment for signatures related to poorly differentiated mPC (basal and NEPC signatures) compared to vehicle, whereas in the matched tumor tissue samples, this enrichment for NEPC-related signatures only became evident at later timepoints, when the tumor was macroscopically growing (29 days) (**Figure 4E**).

In summary, EV-RNAseq recapitulated the expression programs of matched tumors in response to treatment and could detect changes associated to enzalutamide resistance, even earlier than tumor tissue RNAseq.

### Plasma EV-RNAseq in mPC patients undergoing ARSI therapy

We performed longitudinal EV-RNAseq (from samples taken at treatment baseline, after 4 weeks on treatment, and at progression) in mPC patient plasma samples receiving ARSI therapy (n=15 patients, 38 samples), focusing the analysis in MSigDB hallmark and a literature-curated set of gene signatures (**Supplementary Table S2**).

After 4 weeks of therapy, a consistent downregulation of several Hallmark gene sets related to cell proliferation (E2F targets, OXPHOS and mTORC1 signaling), DNA repair (Hallmark DNA repair) and AR signaling (luminal activity and AR-V7) was observed in EV-RNA (**Figure 5A**). In contrast, there was an increase in basal-like and neuroendocrine related gene sets (RB1 loss, NE Labrecque) in EV-RNA upon exposure to ARSI both on treatment (n=13) and at PD (n=9) compared to treatment-baseline samples, in line with known emerging drug resistance mechanisms in PC. Down-regulation of multiple signatures related to AR signaling upon therapy initiation was selectively marked in patients who achieved PSA responses to therapy, whereas the transcriptome of patients who did not respond to therapy remained largely unchanged (**Figure 5B**).

**Figure 5—.**
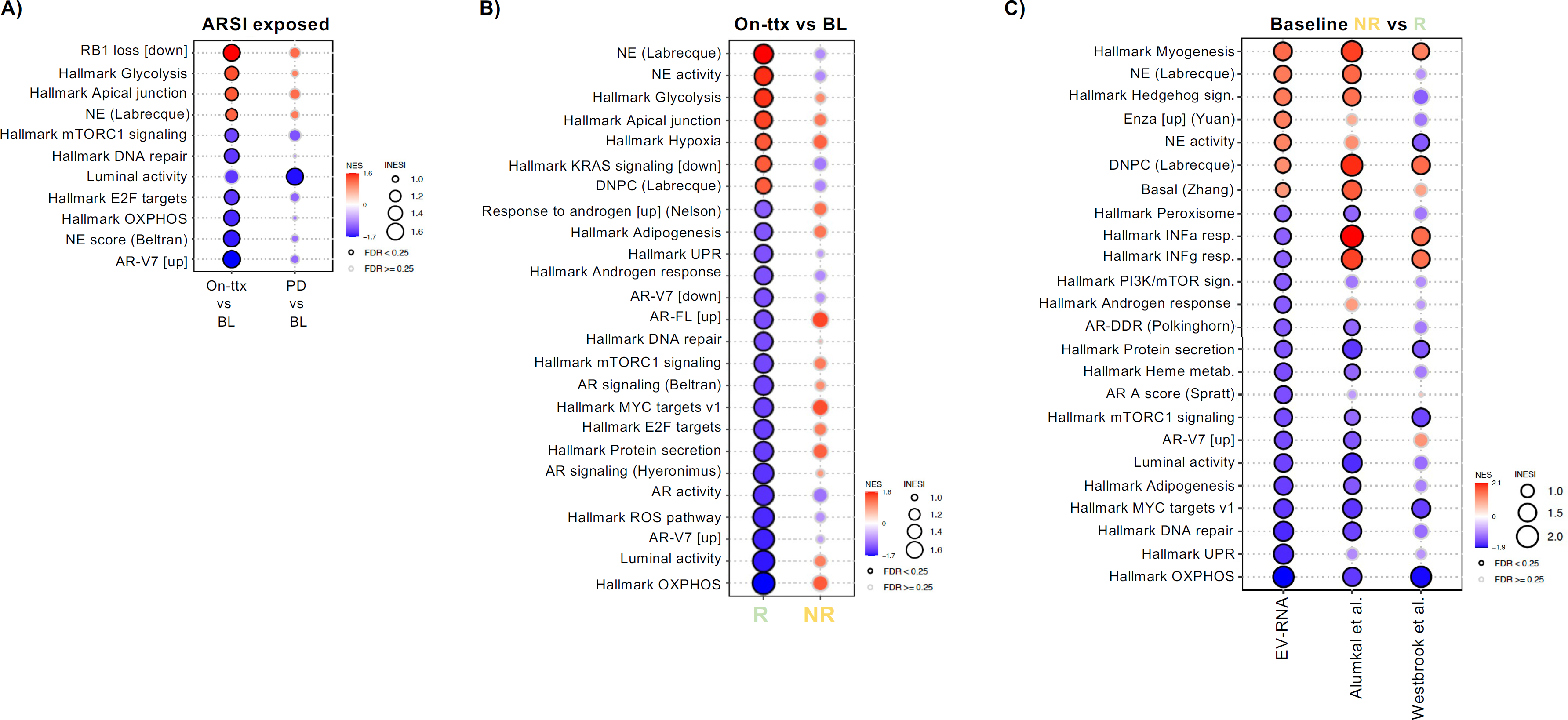
Effects of ARSI treatment on plasma-derived EV-RNA transcriptome and differences between responder and non-responder patients. **a)** Bubble plot showing significant gene set enrichment scores in EV-RNA after treatment with ARSI at 4-weeks of treatment (On-ttx) or at progression (PD) versus baseline (BL). FDR < 0.25 was used for significance. **b)** Bubble plot showing significant gene set enrichment scores in EV-RNA upon ARSI treatment (4 weeks) in responder (R) and non-responder (NR) patients. FDR < 0.25 was used for significance. **c)** Bubble plot showing on the left significant gene set enrichment scores in EV-RNA from non-responder (NR) versus responder (R) patients at baseline; and on the right the validation of these changes in RNA from tumor biopsies of non-responder patients to enzalutamide at baseline (Alumkal et al., and Westbrook et al. cohorts (4,16)). FDR < 0.25 was used for significance. Gene signatures include the MSigDB hallmark collection and literature curated gene sets related with ARSI response and resistance and DNA repair. Normalized enrichment score (NES).

To explore the potential of EV-RNA to predict treatment response in patients, we compared baseline EV-RNA from non-responder (n=6) vs. responder (n=8) patients. Baseline EV-RNA in non-responder patients was enriched for gene sets related to poorly differentiated PC, and showed lower representation of gene sets related to luminal activity, AR signaling, and DNA repair compared to responders (**Figure 5C**). These findings were validated in two previously published cohorts of mPC patients undergoing enzalutamide treatment with available baseline tumor biopsy RNAseq data and clinical outcome data annotated (4,16). Only immune-related signatures were differentially associated with response between tumor biopsies and EV-RNA, probably because of the contribution of the tumor immune infiltrate to bulk tumor RNAseq data. These results demonstrate that EV-RNA recapitulates the transcriptional characteristics of enzalutamide-resistant tumors (AR-low, stemness) in circulation.

## Discussion

In this study, we showed that DNA and RNA contained in circulating EVs secreted by tumor cells reflect PC genomic and transcriptomic features. Critically, we tracked clinically relevant transcriptomic signatures associated with resistance to AR inhibition and lineage plasticity, detecting them early after therapy onset. To the best of our knowledge, this is the first study to deliver an EV-based tumor genomic and whole-transcriptome analysis on clinical samples, expanding the opportunities to study cancer evolution from liquid biopsies.

The field of liquid biopsies in PC has advanced exponentially over the last decade (7,17–19). CtDNA NGS has proven useful for mPC stratification for precision medicine therapies, along with research in other tumor types (20). Nonetheless, resistance to AR inhibition in mPC results from a complex tumor adaptation process involving transcriptomic regulation (5,16). While some genomic backgrounds have been associated with this lineage plasticity, ctDNA NGS seems insufficient to dissect clinical resistances to AR inhibition and to guide therapeutic decisions.

Prior studies demonstrated the presence of tumor mutations in circulating EVs (21–23). In here, we conducted a comparative dual analysis of EV-DNA and matching cfDNA from the same plasma sample. We confirmed the tumor content of circulating EVs by showing that tumor CNAs are represented in EVs from *in vitro* and *in vivo* models, and then in patient samples. In those, EV-DNA CNA profile matched that of same-patient tumor biopsies. Importantly, our data showed that EV-DNA tumor fraction is prognostic in mPC patients.

Previous studies aiming to infer the transcriptional profiles of tumors from liquid biopsies have focused on methylation changes in ctDNA (24,25), fragmentomics (26,27), and more recently, nucleosome positioning (28). In this study, we developed a pipeline for the analysis of mRNA Expression in Circulating EVs (RExCuE) to streamline transcriptomic analysis in liquid biopsies. In contrast to previous studies (23,29–32), this method allowed us to specifically enrich for the protein-coding transcript RNA cargo in EVs. Using this novel approach, we performed transcriptomic profiling of protein-coding genes and detected expression of relevant mPC genes in circulating EV-RNA which have been previously identified in mPC CTCs (15). These tumor-specific features were further supported by our findings when comparing EV-RNA from patients to PBMC from the same patients and EV-RNA from a group of age-matched healthy volunteers. We further characterized circulating EV-RNA using cell-deconvolution protocols, showing an enrichment for epithelial markers in mPC patient-derived EVs.

Recent studies focusing in CTC and ctDNA have shown that early timepoint samples (4 weeks after therapy initiation) can be useful to predict disease outcome (33,34). Here, by comparing longitudinal samples upon therapy exposure, we demonstrated that known gene signatures of therapy response and/or resistance to ARSI (4,5,16,35) can be already detected at such early timepoints in circulation (9 days after treatment in PDX plasma and 4-weeks after treatment in mPC patients). Our data underscores the potential of this minimally invasive biomarker, even more considering the challenges to repeat tissue biopsies longitudinally in clinical practice. This biomarker could assist the development of new combination therapies with AR-targeting agents, particularly those that aim to revert ARSI-resistances (36–38) or are based on cross-talk between pathways at the transcriptional level (39,40). Furthermore, understanding whether tumors remain AR-driven upon clinical progression to ARSI could guide further therapy selection.

Despite the relevance of our findings, this study has some limitations that need to be considered. First, tumor EV-DNA and EV-RNA are subject to dilution among non-tumoral EV in circulation, same as ctDNA is diluted in non-tumoral cell-free DNA (17,34,41,42). In our study, we focused in late-stage prostate cancer patients, where the proportion of tumor related circulating material is expected to be higher. Further studies are needed to assess the sensitivity of our circulating EV-based assays in earlier stages of the disease; other groups are currently investigating alternative isolation methods, based on detecting tumor-specific membrane protein markers in EVs to enrich for tumor-specific EVs, or aiming for single-EV characterization (43,44). Second, our analysis compared EV-DNA to single tumor biopsies but EV-DNA, same as ctDNA, probably encompasses material coming from different metastatic lesions and potentially distinct disease subclones. Our findings have to be interpreted with caution, based on our limited sample size, that however represents the largest analysis of its kind to date.

In summary, this work demonstrates that tumor genomic and transcriptomic profiling of mPC can be studied in circulating EVs. These results highlight the potential of EV analysis to accelerate drug development and to study drug response and resistance mechanisms in PC.

## Methods

### Cell lines and culture

PC cell lines (LNCaP, C4-2, 22Rv1, and PC3) were obtained from LGC standards/ATCC, cultivated according to supplier’s recommendations, STR profiled, and tested regularly for mycoplasma. Cells were cultured in RPMI medium with 10% EV-depleted fetal bovine serum and 1% penicillin-streptomycin.

### Patient and sample characteristics

First, plasma samples from a retrospective cohort of 22 mPC patients were obtained from the Vall d’Hebron University Hospital and used for EV characterization; 12 were used for ultracentrifugation-EVs and 10 for exoRNeasy-EVs characterization. A second set of plasma samples from metastatic castration-resistant PC (n=92) were collected longitudinally during ARSI (abiraterone acetate or enzalutamide) treatment (at baseline, BL; after 4 weeks on treatment, On-ttx; and at progression, PD) from a further cohort of 35 mPC patients. Tissue samples frozen in OCT or FFPE, corresponding to primary or metastatic tumor biopsies from the same patients, were also available for eight patients. Clinical data were retrospectively collected from electronic patient records. All samples and data were obtained after obtaining informed consent to an ethics-approved protocol (Vall d’Hebron Hospital Review Board approval PR(AG)5248).

### Extracellular vesicles purification and characterization

Extracellular vesicles (EVs) from cell line supernatant and plasma used for EV-DNA studies were purified by ultracentrifugation as previously described (10). Cell culture supernatant (*in vitro* experiments) or plasma (*in vivo* and patient samples) underwent serial centrifugation, first at 500g for 10 min, followed by 15 min at 3000g centrifugation and then 20 min at 12,000g. EVs were collected by ultracentrifugation of this supernatant at 100,000g for 70 min, and the pellet was washed in PBS and centrifuged again at 100,000g for 70 min. EVs from cell culture supernatants and plasma used for EV-RNA studies were purified by filtration using the Qiagen exoRNeasy kit. Plasma samples used for EV-RNA isolation with exoRNeasy were pre-filtered (0.2um filter, Millex-GP SLGP033RS, Millipore), following manufacturer instructions, to remove larger vesicles (LVs). When indicated, EVs were further purified by discontinuous density gradient layering each of the 40, 30, 20, 10 and 5% (w/v) iodixanol solutions prepared with Optiprep™ (60%w/v) in 0.25 M sucrose/1 mM EDTA/10 mM Tris-HCl, (pH 7.5). Samples were placed on top and ultracentrifuged at 100,000 x g for 16h at 10 °C. Sequential fractions were collected, washed with PBS and ultracentrifuged at 100,000 x g for 70 min. For comparison purposes, LVs obtained after centrifugation at 12,000g for 20 min (12K pellet) were collected in parallel to iodixanol gradient EV samples.

EVs isolated from cell line models and patient samples were characterized according to the recommendations of the International Society of Extracellular Vesicles (45) based on the expression of small EVs marker proteins (CD9, CD63, and CD81 tetraspanins and TSG101)(46). Furthermore, vesicles were characterized independently by high-resolution imaging of single EV (based on negative stain transmission electron microscopy, TEM), and single particle content and size distribution were analyzed by NTA with Nanosight analysis (NS300 instrument, Malvern Panalytical) equipped with a blue laser (405 nm).

### Negative staining transmission electron microscopy (TEM)

Briefly, the isolated vesicles were fixed in 2% Paraformaldehyde - 0.1 M phosphate buffered saline and applied over carbon-coated copper grids using the glow discharge technique (30 sec, 7,2V, using a Bal-Tec MED 020 Coating System). After additional fixation with 1% glutaraldehyde, the grids were contrasted with 2% uranyl acetate and embedded in methylcellulose. The samples were examined using a transmission electron microscope (FEI Tecnai G2 Spirit, ThermoFisher Scientific, Oregon, USA). Images were taken with a Xarosa digital camera (EMSIS GmbH, Münster, Germany) controlled by Radius software (Version 2.1).

### Plasma processing and DNA extraction

Whole blood was collected in EDTA K2 tubes and processed within 1-3 hours of collection by centrifugation at 500g for 10 min for mouse samples or by centrifugation at 1600g for 10 min for patient samples. The plasma layer was transferred to a new tube and centrifuged at 3000g for 15 min at room temperature (RT). Plasma was stored at -80°C.

Frozen plasma or cell supernatants were used for dual cell-free DNA and EV-DNA purification. The supernatant collected after the first ultracentrifugation was used for cfDNA isolation, whereas the resulting EV pellet was washed, resuspended in PBS, and used for EV-DNA extraction. Cell-free DNA from cell culture supernatants and mouse plasma was manually isolated using the QIAamp MinElute ccfDNA Mini Kit (Qiagen), while cfDNA from patient plasma samples was isolated with QIAsymphony SP using the QIAsymphony DSP Circulating DNA Kit (Qiagen). Pelleted EVs were subjected to DNAse digestion with DNAse I to degrade any cfDNA, followed by proteinase K digestion and DNA isolation with QIAmp DNA Mini Kit (Qiagen), following the manufacturer instructions. Genomic DNA (gDNA) from cultured cells was isolated using the QIAmp DNA Mini Kit (Qiagen), according to the manufacturer protocol. DNA and RNA from patient tumor tissue biopsies and mouse tumors were extracted from 4 sections of 10 µm each from FFPE tumor blocks using the AllPrep® DNA/RNA FFPE kit (Qiagen #80234) according to the manufacturer’s instructions. DNA concentration was assessed using a Qubit 4.0 (Thermo Fisher Scientific). Integrity and fragment size distribution were assessed using a TapeStation 4200 (Agilent Technologies).

### DNA libraries, sequencing and data processing

gDNA from cultured cells and tumors and EV-DNA were subjected to mechanical fragmentation using a Covaris M220 ultrasonicator. Libraries for gDNA, cfDNA, and EV-DNA samples were constructed using the KAPA HyperPrep kit (Roche), following the manufacturer’s instructions. Libraries were sequenced on a HiSeqX to an average depth of 0.5X with 150 bp paired-end reads per sample.

Raw fastq files were processed as follows: (1) trimmomatic v0.39 was used to remove adapters; (2) reads were mapped to the hg19 human genome with Bowtie2 v2.3.5.1; (3) duplicates were removed with samtools; (4) segmentation was performed with ReadCounter with a 500Kb window, removing low-quality reads (<Q20); (5) the segmentation file was the input of the ichorCNA package in R (19), which calculated segment-based copy number, tumor fraction, and ploidy. IchorCNA lower limit of sensitivity for detecting the presence of tumor in circulation is 3%; hence, we considered “EV-DNA detectable” all those samples with a tumor fraction ≥ 4%.

## RNA isolation

RNA from FFPE tumors (xenografts and patient biopsies) was extracted using an AllPrep® DNA/RNA FFPE kit (Qiagen). The AllPrep® DNA/RNA/miRNA kit (Qiagen) was used to isolate RNA from tumor OCT blocks. Fresh-frozen mouse tumors were mechanically disrupted using TissueLyser II (Qiagen), and RNA was isolated using the RNeasy kit (Qiagen) following the manufacturer’s instructions.

EV-RNA from mouse and patient plasma was extracted using the ExoRNeasy kit (Qiagen), following the manufacturer’s instructions. The extracted material was DNAse treated using the RNase-free DNAse set from Qiagen. The RNA concentration was assessed using a Nanodrop spectrophotometer. Integrity and fragment size distribution were assessed on a TapeStation 4200 using a High-Sensitivity RNA ScreenTape (Agilent Technologies).

### RExCuE RNA libraries, sequencing and data processing

The RExCuE protocol for RNAseq library preparation was developed *in house* as a modification of the SmartSeq2 protocol (47) to generate poly(A)-captured libraries. Refined reverse transcription and template switching were performed using the SuperScript IV reverse transcriptase protocol (Invitrogen) with a modification of the SmartSeq 2 protocol. After reverse transcription, PCR pre-amplification was carried out and the PCR product was purified using a 1:1 ratio of SPRI beads (Clean NA). The pre-amplified product size distribution was checked on a High-Sensitivity D5000 ScreenTape (Agilent) and quantified on a Qubit 4.0. The pre-amplified cDNA was mechanically fragmented using a Covaris M220 ultrasonicator and used as input for RNA libraries using the KAPA HyperPrep kit (Roche) following the manufacturer’s instructions, with modifications in both the universal adaptor concentration and amplification cycles according to the KAPA-HyperPrep manual KR0961-v6.17.

Libraries were sequenced on an Illumina NovaSeq, PE150 for an average of 50 million reads/sample. Raw fastq files were processed as follows: (1) BBDuk was used to identify and remove possible rRNA contamination; (2) Trim-Galore 0.6.7 was used to remove adapters; (3) Reads were mapped to the GRCh38 human genome with STAR 2.7.9a. Gene expression levels were quantified with featureCounts from Subread 2.0.3 (48). Gene expression was transformed to logCPM and normalized using the quantile method with voom from the limma package (49) in R. Genes with little variation across samples (sd ≤ 0.01) were removed. Non-protein-coding genes were removed from the analysis. We applied RemoveBatchEffect to adjust the expression data according to batch effect. For mouse RNA and EV-RNA, reads were separately mapped to the GRCh38 human genome and GRCm39 mouse genome with STAR 2.7.9a; reads of human origin were isolated with XenofilteR (50). An average of 1 million mapped human reads were available for gene expression analysis after mouse read removal in LNCaP and C4-2 xenograft’s plasma and 64 and 11 million mapped human reads were obtained after mouse removal for LNCaP and C4-2 xenograft tumor RNA, respectively; while an average of 9 million and 40 million mapped human reads were available for analysis after mouse read removal in plasma EV-RNA and tumor RNA from PDXs, respectively. Gene expression levels were quantified using feature counts; gene expression was transformed and filtered as described above.

### Differential expression and GSEA analysis

To identify differentially expressed genes (DEGs), we used the Empirical Bayes linear modeling framework from the limma package (49) in R. The DEGs were defined by a |log2FC| > 1 + FDR-adjusted P < 0.05. Gene set enrichment analysis was performed using the fgsea package in R. As a source for functional annotation, we used the Hallmark gene set from MSigDB and a collection of gene sets extracted from the literature (16,35,51–63)(**Supplementary Table S2**). Nominal P-value< 0.05 and FDR<0.25 were used as cutoffs for significance. Functional enrichment analysis was performed with fgseaMultilevel from the FGSEA R package.

### Gene Ontology (GO) analysis

Statistically significantly upregulated ensemble gene IDs from our DEG analysis (FDR<0.05) were assessed for enrichment in the PANTHER classification system (v.17.0). GO biological process complete database was used for functional classification.

### Cell deconvolution analysis

Raw counts data was transformed to transcripts per million counts (TPM) to perform deconvolution analyses. Deconvolution analyses were performed in R using xCell package (v1.1.0)(12).

### Enzymatic protection assays

Plasma-derived EVs, isolated by ultracentrifugation, or cfDNA samples were subjected to DNAse digestion by treating them with 1 µl of DNAse I (Thermofisher) in DNAse digestion buffer (1×) and incubated at RT for 15 min, followed by enzyme inactivation at 70 °C for 10 min. After digestion, EVs were incubated with Proteinase K and lysed to perform DNA extraction using the QIAmp DNA Mini Kit (Qiagen) according to the manufacturer’s protocol. Plasma-derived EVs, isolated with the exoRNeasy kit (Qiagen), were subjected to RNAse digestion by treating them with an RNAse A/T1 mix (Thermo Fisher) and incubating them at 30 °C for 20 min followed by digestion with Proteinase K at 56 °C for 5 min. The integrity and fragment size distribution for both EV-DNA and RNA were assessed using TapeStation 4200.

### Western Blotting

SDS-PAGE (10% reducing, 5% 2-β mercaptoethanol) was used for protein expression analysis. The following antibodies were used: CD63 (H5C6; #556019; BD Biosciences), CD81 (B11; sc-166029; Santa Cruz Biotechnology), CD9 (MM2/57; CBL-162; Millipore), TSG101 (C2; sc-7694; Santa Cruz Biotechnology), β-actin (C4; sc-47778; Santa Cruz Biotechnology), and GAPDH (ab9485; Abcam). Secondary antibodies conjugated to horseradish peroxidase (goat anti-mouse and anti-rabbit IgG; ab97040 and ab205718, Abcam, Cambridge, UK) were used. SuperSignal West Dura Extended Duration Western Blotting Substrate was used and visualized on Amersham Imagen 600 (GE).

### Cell-lines derived xenografts

For the 22Rv1 xenografts, 2 million cells resuspended in Matrigel (Corning) were injected subcutaneously into both flanks of NMRI-Foxn1 nu/nu (n=3). Tumors were measured using calipers, and body weight was monitored weekly. Animals were sacrificed by cervical dislocation under general anesthesia when the tumor volume exceeded 2000mm^3^. Whole blood was collected by heart puncture in EDTA tubes and processed fresh. Plasma from three mice was pooled and frozen at -80°C for downstream analysis. Tumors were collected, snap-frozen or fixed overnight in formalin, and embedded in paraffin.

For LNCaP xenografts, 4 million cells resuspended in Matrigel (Corning) were injected subcutaneously into both flanks of castrated NMRI-Foxn1 nu/nu mice (n=9). Tumors were measured weekly using calipers and body weight was monitored weekly. Once the animals reached a tumor volume of 600mm^3^, they were randomized to either vehicle (n=3) or enzalutamide (n=6) treatment. Enzalutamide was administered orally (10 mg/kg every day, 6 days on-1 day off). For C4-2 xenografts, 3 million cells resuspended in Matrigel (Corning) were injected subcutaneously into both flanks of castrated NMRI-Foxn1 nu/nu mice (n=8). Six-week-old NMRI-Foxn1 nu/nu mice were castrated surgically. C4-2 cells were injected 2-weeks after the surgery (8 week-old mice). Tumors were measured weekly using calipers and body weight was monitored weekly. Once the animals reached a tumor volume of 700mm^3^, they were randomized to either vehicle (n=3) or enzalutamide (n=5) treatment. Enzalutamide was administered orally (10 mg/kg every day, 5 days on-2 days off).

The animals were sacrificed by cervical dislocation under general anesthesia. Whole blood was collected by heart puncture in EDTA tubes and processed fresh. Plasma from each mouse was individually frozen at -80 °C. Tumors were collected, weighed, snap-frozen or fixed overnight in formalin, and embedded in paraffin blocks.

All experimental protocols were approved and monitored by the Vall d’Hebron Institute of Research Animal Experimentation Ethics Committee (CEEA; registration number 68/20) in accordance with relevant local and EU regulations.

### Patient-derived xenograft (PDX)

Patient-derived xenograft P886 was generated from a transrectal prostate biopsy of a patient with *de novo* metastatic hormone-naïve PC. Briefly, one tumor biopsy core was implanted with growth factor-enriched Matrigel (Corning) subcutaneously in the flank of an NSG NOD SCID gamma mouse. Tumors were collected when they reached a volume of 600mm^3^ and serially passaged into both flanks of NSG NOD SCID gamma mice.

Tumors from PDX P886 were collected and digested with a mixture of collagenase type II and TryplE to generate a cell suspension. Five hundred thousand cells were injected subcutaneously into each flank of NSG NOD SCID gamma mice. Tumors were measured twice weekly using calipers and body weight was monitored three times per week. Animals were randomized when tumors reached 700 mm^3^ into two treatment arms: vehicle (n=2) or enzalutamide (n=4). Enzalutamide was administered orally (30 mg/kg) every day (6 days on, 1 day off) for 9 days or until the mice progressed to treatment (29 days). The animals were sacrificed by heart puncture under general anesthesia.

Whole blood was collected by heart puncture in EDTA tubes and processed fresh. Plasma was frozen at -80 °C. The tumors were excised, weighed, frozen in OCT blocks, or fixed in formalin overnight, and embedded in paraffin.

### Histological and immunohistochemistry analysis

Hematoxylin and eosin staining was performed on the FFPE sections. Ki67 and AR IHC were performed using the Discovery Ultra Autostainer (Ventana Medical Systems, Tucson AZ) and the following antibodies: Androgen Receptor Rabbit Monoclonal Antibody, clone SP-107 (760-4605, Roche); Ki67 mouse monoclonal antibody, clone MIB-1 (M7240, Dako).

### Statistical analysis

Statistical analysis and graphs were performed using GraphPad Prism 8 (version 8.4.0) and the R package (Version 4.1.2). Error bars in graphs represent mean ± s.e.m. statistical significance was determined using two-tailed Student’s t-test or one-way ANOVA. Correlations were determined using Pearson’s correlation, and Pearson’s correlation coefficient (r) was used to measure the strength of the association between the two variables analyzed.

For the clinical outcome correlative analysis, the following endpoints were considered: PSA response (PSA50, defined as a decline in PSA>=50% compared to baseline) was used to classify patients into responders and non-responders; progression-free survival was defined as time from treatment initiation to radiographic progression, unequivocal clinical progression leading to a change in treatment, or death. Univariate progression-free survival analysis was performed using the Kaplan-Meier estimator (log-rank test). Statistical significance was set at P < 0.05.

## Data availability

All sequencing data generated in this study has been deposited at NCBI GEO repository with accession number GSE221709.

