## Supplemental Figures for "Circulating tumor extracellular vesicles to monitor metastatic prostate cancer genomics and transcriptomic evolution"

**Supplementary Figure 1— a)** Immunoblot analysis for sEV markers (TSG10, CD9 and CD81) in LNCaP, C4-2, 22Rv1 and PC3 prostate cell lines. Beta-actin is shown as an internal loading control. **b)** Transmission electron microscopy (TEM) of LNCaP, C4-2, 22Rv1 and PC3 EVs. Scale bar, 200 nm. **c)** EV size (median, nm) and number were evaluated by NanoSight particle tracking in LNCaP, C4-2, 22Rv1 and PC3 EVs. **d)** Analysis of sEV sequential fractions obtained after iodixanol density gradient of a 22Rv1-derived sEV preparation. Quantification of dsDNA (ng) is shown for LVs (12K) and each of the collected fractions (F1-12). Markers for sEV (CD81 and TSG101) were assessed in F1-F12. **e)** Representation of whole genome CNA profile for Tumor DNA and EV-DNA from LNCaP, C4-2, 22Rv1 and PC3 cell lines and tumor DNA and plasma-derived EV-DNA from 22Rv1 mouse xenograft. Amplifications are depicted in red and deletions in blue. Similarity between CNA in DNA from tumor cells (TumDNA) and EV-DNA is given by Pearson correlation score ( r ); in all cases, P-value<0.0001 for matched TumDNA-EV-DNA pairs.

**Supplementary Figure 2— a)** Immunoblot analysis for sEV markers (TSG101, CD9 and CD63) in plasma circulating EVs from mPC patients (n=4). GAPDH is shown as an internal loading control. **b)** Transmission electron microscopy (TEM) of plasma circulating EVs from mPC patients (n=3). Scale bar, 200 nm. **c)** EV size (median, nm) and number were evaluated by NanoSight particle tracking in plasma circulating EVs from mPC patients (n=3).

**Supplementary Figure 3— a)** Violin plots showing the concentration (ng/mL plasma) of matching circulating EV-DNA and cfDNA longitudinally collected at baseline (BL), after 4-weeks of treatment (On-ttx) or at disease progression (PD) from mPC patients (n=35). Table below indicates mean concentration and 95% CI per timepoint. **b)** Stacked bar chart showing the distribution of patients with a detectable TF by sWGS in both cfDNA and EV-DNA, only in one of them (EV-DNA only or cfDNA only) or undetected. Table below indicates mean TF and 95% CI per timepoint. **c)** Scatter plots representing the correspondence between TF in EV-DNA (y-axis) and their matching cfDNA (x-axis) in longitudinally collected samples from mPC patients. Each dot represents one patient. Similarity in gene expression between EV-DNA and cfDNA TF is given by Pearson correlation score ( r ) and P-value. **d)** Table summarizing clinical attributes and their association with time to progression. P-values and Hazard Ratio shown were determined using the Cox proportional hazards model.

**Supplementary Figure 4— a)** EV size (median, nm) and number distribution by NanoSight particle tracking analysis of plasma-derived EVs isolated with exoRNeasy kit from mPC patients (n=2). **b)** Upper, immunoblot analysis for sEV markers (TSG101, CD9 and CD81) of plasma-derived EVs isolated with exoRNeasy kit from mPC patients (n=3). GAPDH is shown as an internal loading control. Lower, transmission electron microscopy (TEM) of plasma-derived EVs isolated with exoRNeasy kit from mPC patients (n=3). **c)** Analysis of sEV sequential fractions obtained after iodixanol density gradient of a ExoRNeasy-derived plasma sEV preparation. Quantification of EV-RNA (ng) is shown for each of the collected fractions (F1-12). Markers for sEV (CD9 and TSG101) were assessed in F1-F12. **d)** RNAse digestion assay of EV-RNA. Tapestation quantification of RNA concentration (pg/µl) from untreated (white) and RNAsetreated (checkered) samples of plasma-derived EVs (n=4). **e)** Bar graph representing the percentage of RNA biotypes detected in EV-RNA sequencing following our optimized library preparation (see Methods).

**Supplementary Figure 5— a)** Gene Set Enrichment analysis of DE genes in mPC EV-RNA vs HV EV-RNA using GO (gene ontology). **b)** Violin plots showing distribution of individual enrichment scores for the significantly different cell subtypes in healthy volunteers and mPC patients' EV-RNA: immature dendritic cells (iDCs), keratinocytes and neurons. **c)** Cell type enrichment analysis in RNAseq data from matching tumor biopsies (TBx RNA) (n=8). Individual patient enrichment scores are shown in the heatmap.

**Supplementary Figure 6— a)** Representative H&E and immunohistochemical staining for Ki67 and AR for PDX tumors treated with Enzalutamide (Enza1 and Enza2) for 9 and 29 days or Vehicle (VH1 and VH2). **b)** Genomic characterization of PDX P886. Representation of whole genome CNA profile for tumor DNA from PDX P886 (TumDNA) and plasma-derived cfDNA and EV-DNA. Amplifications are depicted in red and deletions in blue. Similarity between CNA is given by Pearson correlation score ( r ); P-value<0.0001. **c)** Size-distribution of RNA fragments in circulating EV-RNA collected from PDXs P886 plasma. Scatter plot showing the percentage of fragments of different sizes found in EV-RNA.

#### SUPPLEMENTARY FIGURE 1

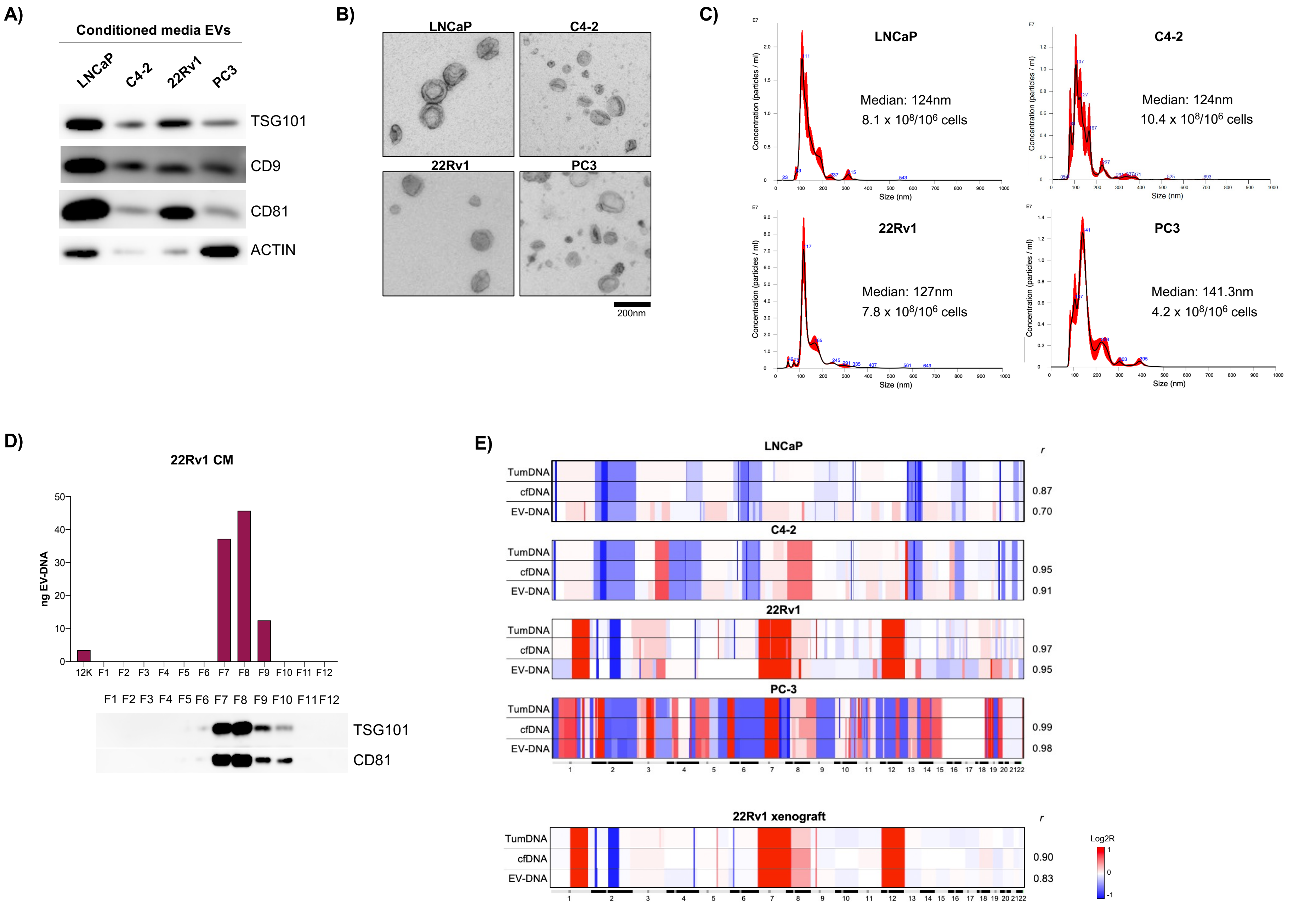

**SUPPLEMENTARY FIGURE 2**

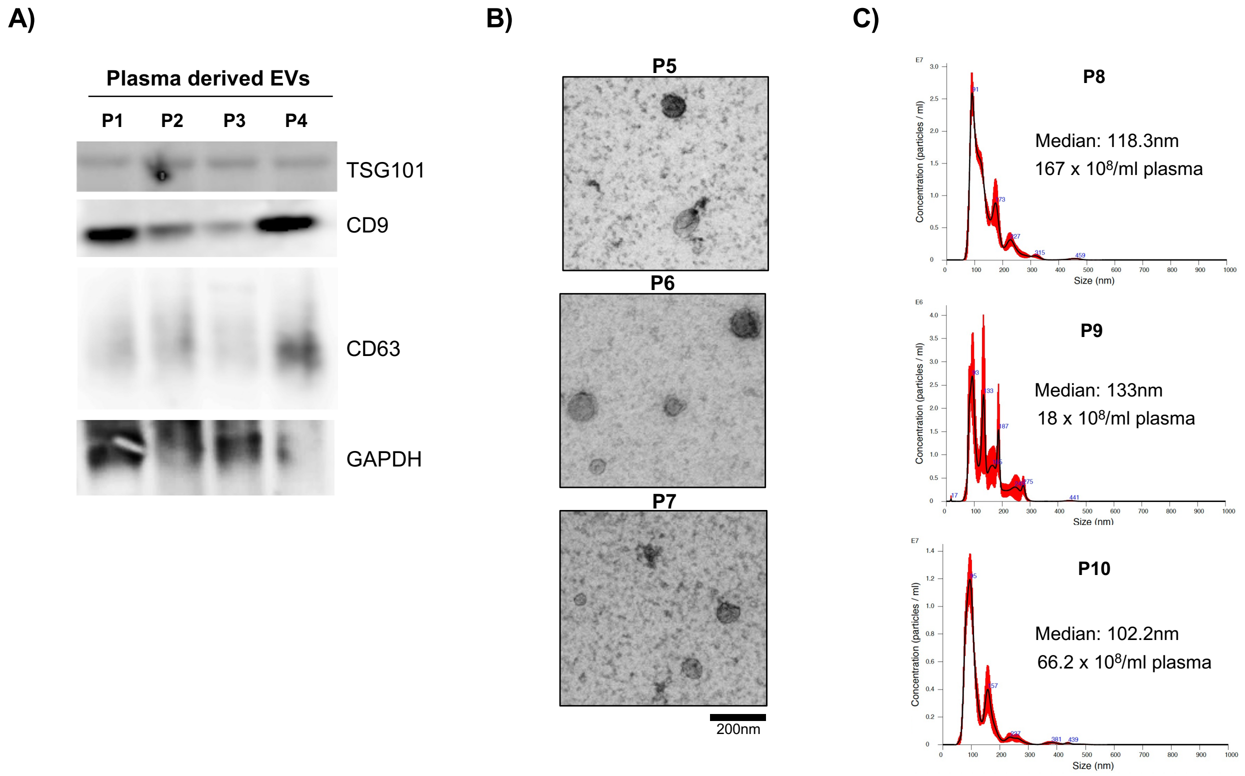

### SUPPLEMENTARY FIGURE 3

## A)

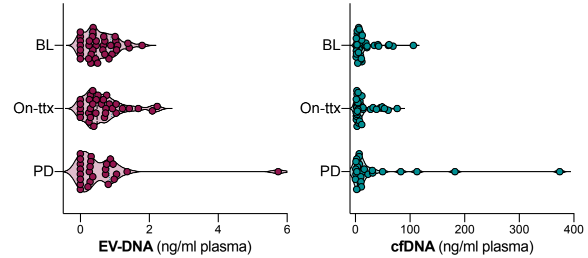


| | EV-DNA (ng/ml) Mean (95% CI) | cfDNA (ng/ml) Mean (95% CI) |
|---|---|---|
| Baseline (BL) | 0.56 (0.39-0.71) | 18.4 (10.3-26.5) |
| On treatment (On-ttx) | 0.64 (0.43-0.85) | 16.2 (8.9-23.5) |
| Progression (PD) | 0.64 (0.17-1.1) | 39.7 (6.1-73.4) |
| All timepoints (n=92) | 0.61 (0.46-0.76) | 23.7 (14-33.4) |

## B)

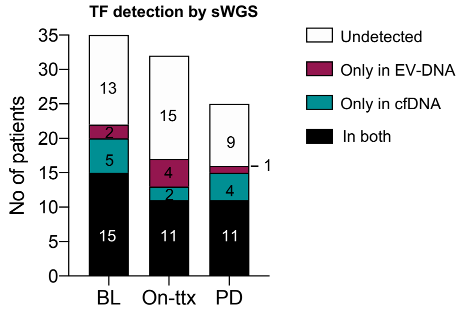


| | EV-DNA TF (%) Mean (95% CI) | cfDNA TF (%) Mean (95% CI) |
|---|---|---|
| Baseline (BL) | 11.3 (5.5-17.2) | 12.9 (7.6-18.3) |
| On treatment (On-ttx) | 9.2 (4.6-13.8) | 11.3 (6.0-16.7) |
| Progression (PD) | 14.3 (5.7-22.9) | 16.4 (8.0-24.7) |
| All timepoints (n=92) | 11.4 (7.9-14.9) | 13.3 (9.9-16.8) |

## C)

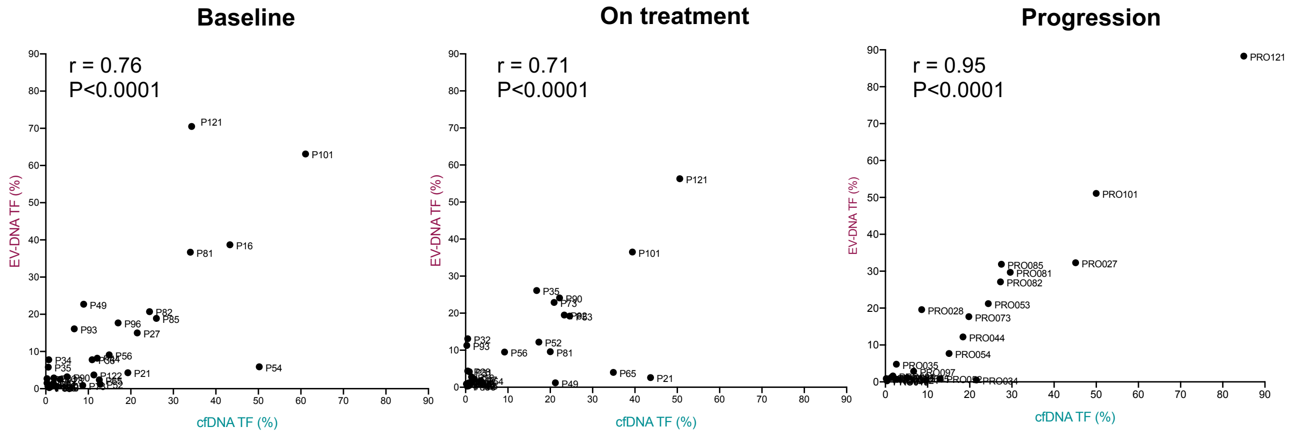


## D)

| | No. Patients | Time to progression: Median (months) | HR (95% CI) | P value |
|---|---|---|---|---|
| **EV-DNA** | | | | |
| Detectable (TF≥4%) | 17 | 11 | 2.29 (1.02-5.12) | **0.044** |
| Undetectable (TF<4%) | 18 | 16 | | |
| **PSA** | | | | |
| ≥10 ng/ml | 25 | 11 | 4.37 (1.59-11.99) | **0.004** |
| <10 ng/ml | 10 | 24 | | |
| **LDH** | | | | |
| ≥250 UI/L | 22 | 12 | 1.81 (0.71-4.61) | 0.213 |
| <250 UI/L | 11 | 16 | | |
| **ALP** | | | | |
| ≥147 UI7L | 11 | 8 | 2.59 (1.11-6.04) | **0.028** |
| <147 UI/L | 24 | 14 | | |

**SUPPLEMENTARY FIGURE 4**

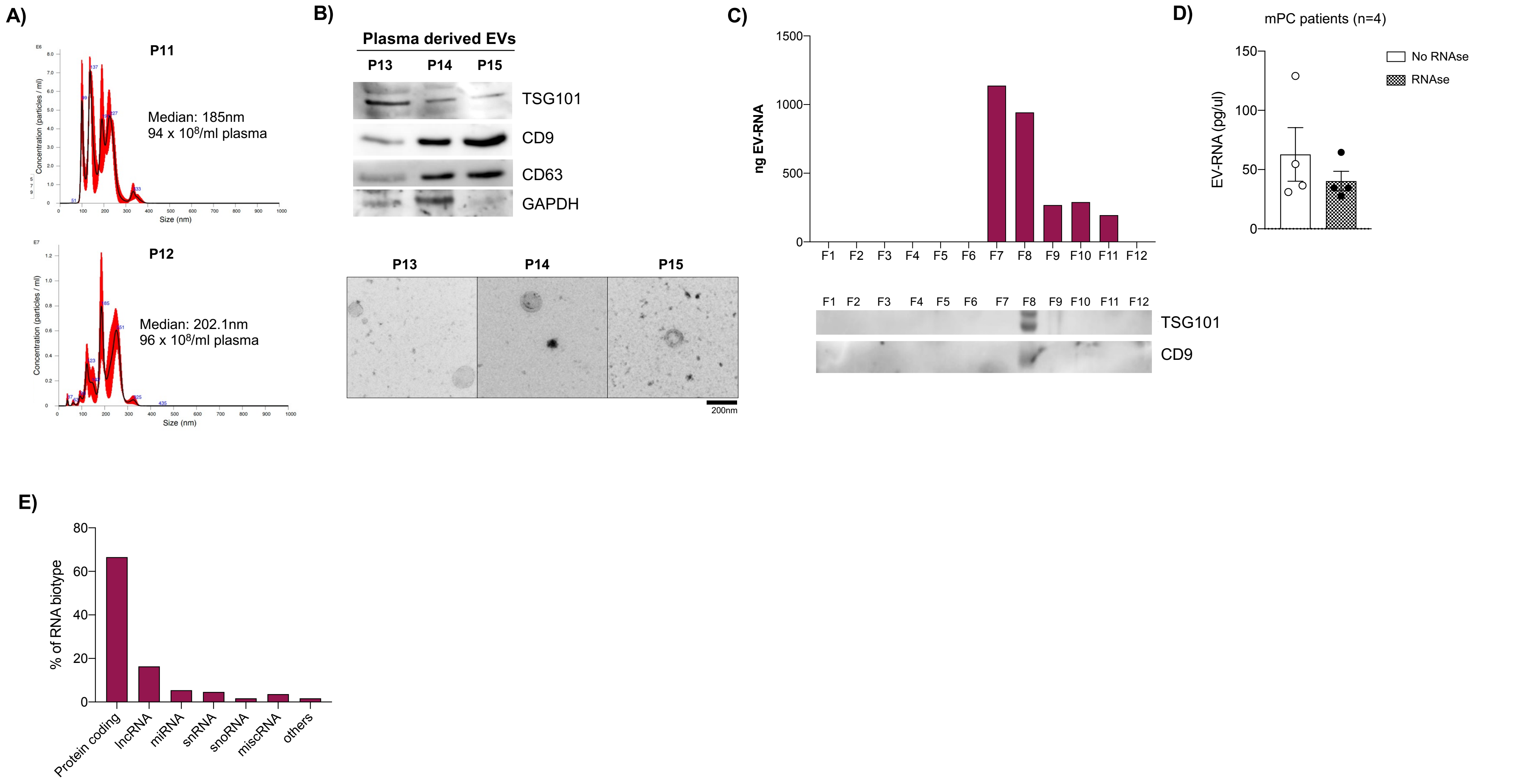

### SUPPLEMENTARY FIGURE 5

A)

| GO term Id | GO term Definition | FDR |
|---|---|---|
| **GO:0009888** | tissue development | 2.600E-02 |
| GO:0060429 | epithelium development | 4.070E-02 |
| GO:0030855 | epithelial cell differentiation | 2.244E-04 |
| GO:0009913 | epidermal cell differentiation | 2.500E-09 |
| GO:0030216 | keratinocyte differentiation | 2.500E-09 |
| GO:0031424 | keratinization | 1.000E-10 |
| GO:0008544 | epidermis development | 1.25E-08 |
| GO:0009913 | epidermal cell differentiation | 2.500E-09 |
| GO:0030216 | keratinocyte differentiation | 2.500E-09 |
| GO:0031424 | keratinization | 1.000E-10 |

B)

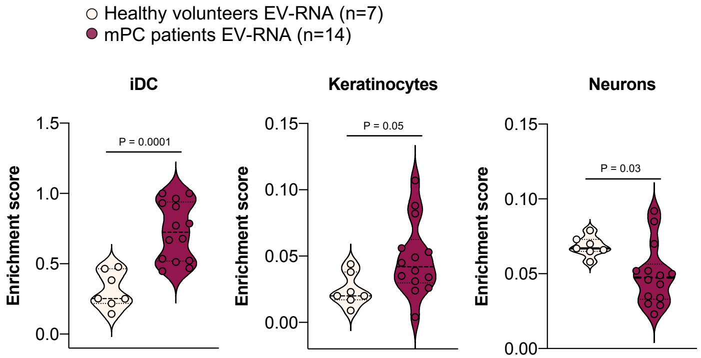


C)

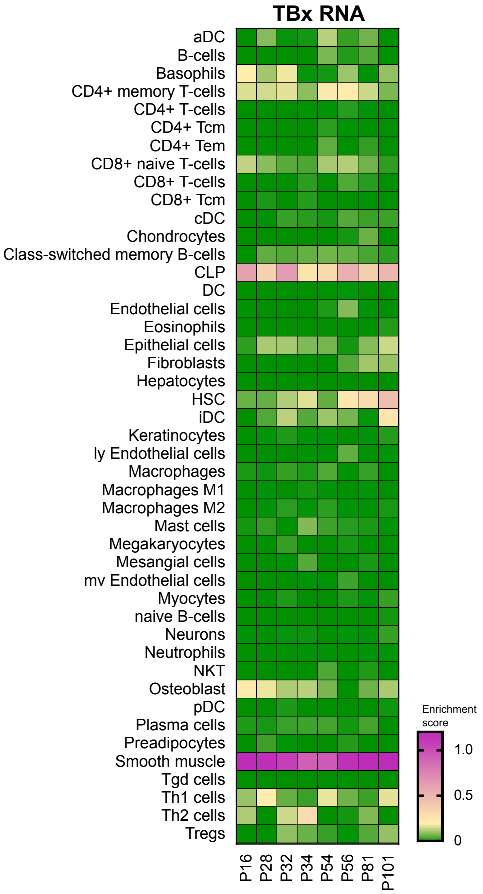

#### SUPPLEMENTARY FIGURE 6

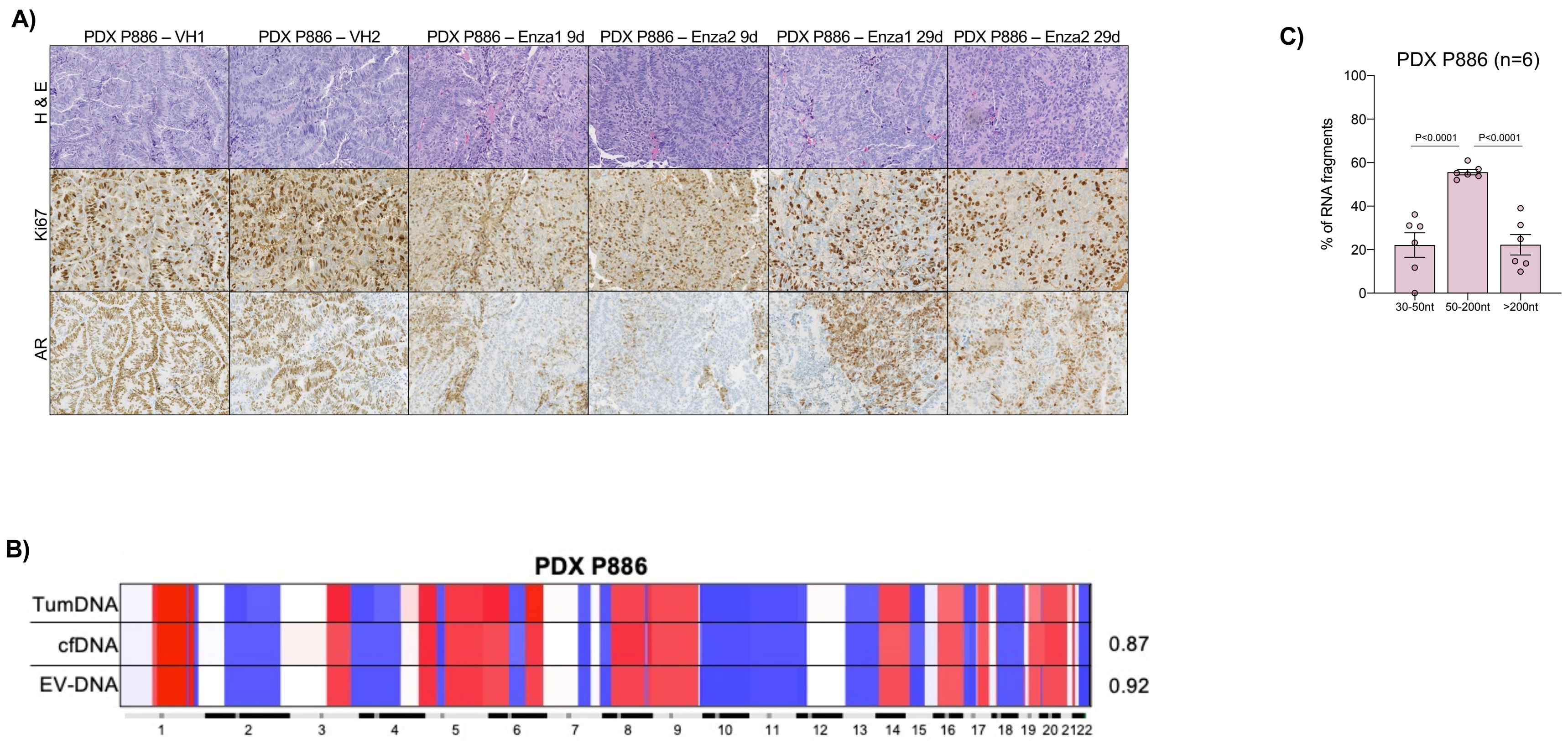
